# A communication hub for phosphoregulation of kinetochore-microtubule attachment

**DOI:** 10.1101/2023.11.22.568383

**Authors:** Jacob A. Zahm, Stephen C. Harrison

## Abstract

The Mps1 and Aurora B kinases regulate and monitor kinetochore attachment to spindle microtubules during cell division, ultimately ensuring accurate chromosome segregation. In yeast, the critical attachment components are the Ndc80 and Dam1 complexes (Ndc80c and DASH/Dam1c, respectively). Ndc80c is an 600-Å long heterotetramer that binds microtubules through a globular “head” at one end and centromere-proximal kinetochore components through a globular knob at the other end. Dam1c is a heterodecamer that forms a ring of 16-17 protomers around the shaft of the single kinetochore microtubule in point-centromere yeast. The ring coordinates the approximately eight Ndc80c rods per kinetochore. In published work, we showed that a site on the globular “head” of Ndc80c, including residues from both Ndc80 and Nuf2, binds a bipartite segment in the long, C-terminal extension of Dam1. Results reported here show, both by in vitro binding experiments and by crystal structure determination, that the same site binds a conserved segment in the long N-terminal extension of Mps1 and a similarly conserved segment in the N-terminal extension of Ipl1 (yeast Aurora B). Together with results from experiments in yeast cells and from biochemical assays reported in two accompanying papers, the structures and graded affinities identify a communication hub for ensuring uniform bipolar attachment and for signaling anaphase onset.

## Introduction

Faithful chromosome segregation in cell division requires bipolar spindle attachment of all sister chromatid pairs. Tension between two sister kinetochores signals biorientation, as the chromosome arms are still cohesin-linked [1–3]. Multiple phosphorylation and dephosphorylation events, in which the protein kinases Mps1 and Aurora B (Ipl1 in *S. cerevisiae*) are instrumental [4–18] transduce the signals reporting attachment/detachment or tension/no-tension, deferring anaphase onset until all kinetochore pairs have attached to opposite spindle poles (the “spindle assembly checkpoint”: SAC, reviewed in [19]). These kinases have interrelated and apparently redundant roles in the error correction needed to achieve uniform biorientation; that is, they participate actively in generating correct attachments, by modifying various kinetochore components, rather than simply reporting attached and unattached states [1, 9, 17, 20, 21]. Their activities are also important at stages of cell division other than the assembly of an anaphase-competent mitotic apparatus, adding further complexity to analysis of their specific functions.

The components of the spindle microtubule attachment machinery in yeast are the Ndc80 and Dam1 complexes (Ndc80c and DASH/Dam1c, respectively). Ndc80c, a heterotetramer of Ndc80, Nuf2, Spc24, and Spc25, is a 600 Å long rod, largely two-chain coiled-coil, with an Ndc80/Nuf2 globular “head” at one end and an Spc24/Spc25 globular knob at the other [22, 23]. The Ndc80 and Nuf2 globular regions are both calponin-homology (CH) domains [24, 25]. Together with an approximately 100-residue, N-terminal extension, the Ndc80 CH domain forms the principal microtubule contact [26, 27]. DASH/Dam1c is a heterodecamer, about 17 copies of which form a ring around the microtubule shaft at the kinetochore (plus) end [28–31]. Through contacts from C-terminal extensions of four Dam1c components, Dam1, Ask1, and Spc19/Spc34, the ring organizes the multiple Ndc80 rods (estimated to be about eight: [32]) surrounding the single kinetochore microtubule [31, 33].

Major targets of Mps1 are a set of so-called MELT repeats on Spc105/Knl1, components of an intermediate, “adaptor” complex that links Ndc80c with centromere-proximal assemblies, which in turn recruit the SAC signaling components [34, 35]. Loss of Mps1 activity leads to prompt dephosphorylation of Spc105/Knl1 and therefore loss of a contribution to the SAC from the kinetochore in question [4, 36]. Mps1 associates with the globular, Ndc80:Nuf2 head of Ndc80c, the main microtubule (MT) attachment component of kinetochores in nearly all eukaryotes [37, 38]. Studies of the metazoan kinase have suggested that MTs and Mps1 compete for Ndc80c binding, leading to the view that binding of Mps1 reports an unattached Ndc80c and that attachment ejects it [37, 38]. Experiments with isolated yeast kinetochores indicate that autophosphorylation of Mps1 may be sufficient to release it from Ndc80c [39], and an accompanying paper [40] shows directly that yeast Mps1 occpancy is fully compatible with microtubule binding, while also suggesting that autophosphorylation alone may not fully explain the dynamics of Mps1 in cells. The difference between the two sets of observations could reflect a difference between metazoan and yeast Mps1, as additional contacts with Ndc80 may be present in the former [38] involving an Mps1 N-terminal segment that has no homologous representation in the latter (see Discussion).

Substrates of Ipl1 in yeast include the Ndc80c (in particular, the N-terminal extension of Ndc80) and the DASH/Dam1 complex (Dam1c) [8, 10, 17, 41, 42]. Phosphorylation of certain Ser/Thr residues in these complexes weakens microtubule attachment. Absence of tension presumably favors Ipl1 access to these sites, thereby promoting error correction. In the case of syntely, for example, in which both sister chromatids are attached to the same pole, phosphorylation would weaken the contact, ultimately freeing one or both of the sister kinetochores to “try again” and reconnect correctly.

Ipl1/AuroraB is the catalytic component of the heterotetrameric chromosome passenger complex (CPC)[43]. A long, potentially extended connector, Sli15/INCENP, links Ipl1 at one end with two regulatory subunits, Bir1/Survivin and Nbl1/Borealin, at the other end. Bir1 dependent association of the CPC with centromeres during the onset of mitosis has led to the suggestion that the length of the INCENP “tether” could determine the likelihood of Ndc80 phosphorylation and that an increase under tension of the distance between centromeric chromatin and the Ipl1 sites on the Ndc80 N-terminal extension could be part of the tension-sensing mechanism [44]. Some studies in yeast have questioned this picture of a distance gradient, however, and an alternative model postulates local, tension-induced changes at the Ndc80-MT interface that sequester the Ipl1 target sites or reduce the frequency of their dissociation [45, 46].

Sorting out the interrelated functions of Mps1 and Ipl1 clearly requires direct structural pictures of their association with kinetochore components. We show here, biochemically and structurally, that both Mps1 and Ipl1 bind the site on the Ndc80/Nuf2 head of Ndc80c that we have shown previously to bind an Ipl1-phosphoregulated peptide segment near the C-terminus of Dam1 [47]. The segment of Mps1 that associates with Ndc80c is in the middle of the ~430-residue N-terminal extension of the kinase domain; the associating segment of Ipl1 is in a much shorter N-terminal extension of the kinase domain. The binding motifs are conserved in their key features across point-centromere yeast.

In related work, Pleuger et al [40] and Parnell et al [48] have characterized Mps1:Ndc80c interactions both in vivo and in vitro; their analyses of mutant phenotypes together show the functional relevance of contacts in the Mps1:Ndc80c structure reported here. The common binding surface for Dam1, Mps1, and Ipl1 on the Ndc80c head is thus a central communication point for establishing the SAC, for triggering error correction and ultimately silencing the SAC, and for generating robust, bipolar attachments as the outcome.

## Results

### Mps1 interaction with Ndc80c

The long, N-terminal extension (residues 1-430) of yeast Mps1 has no immediately evident homology with sequences in metazoan Mps1 shown to interact with Ndc80c. Genetic analysis has suggested that residues 151-200 function in kinetochore biorientation, while residues 201-300 function in spindle-pole body duplication [49]. We examined conservation among the yeast orthologs by iterative searches with Jackhmmer [50]. Using the N-terminal 430 residues of *S. cerevisiae* Mps1 as query, we identified five segments conserved to one degree or another among point-centromere yeasts (Fig. 1A). We carried out a set of pulldown experiments, expressing them as GST-fusions and incubating them with the “dwarf” Ndc80c construct (Ndc80c^dwarf^)[51] immobilized on Ni-NTA agarose (Fig. 1B). In this format, we expected avidity from GST dimerization and multivalent display of Ndc80c to enhance the assay sensitivity. Ndc80c captured only the GST fusion of a peptide containing residues 151-171 (Fig. 1B, lane 12), which included the most conserved region with a consensus sequence (151)RRxRRF(I/L/F)(3-4x)RxxxLGPAxR(170)(Fig. 1C). Lysine appears occasionally in place of arginine, except for the invariant R at the end of the motif. The consensus is an essentially bipartite motif, with a set of basic residues and two hydrophobic residues at one end connected somewhat variably to GPAxR at the other. A multi-chain Alphafold2 (AF2) prediction [52] in which the input was Mps1 residues 150-200, Ndc80 residues 114-335, and Nuf2 residues 1-173 showed an interaction that included the consensus motif (Fig. S1A). We used this prediction to restrict the length of the peptide used for binding experiments, as described in the next section, and to design the construct that yielded a crystal structure.

**Figure 1.**
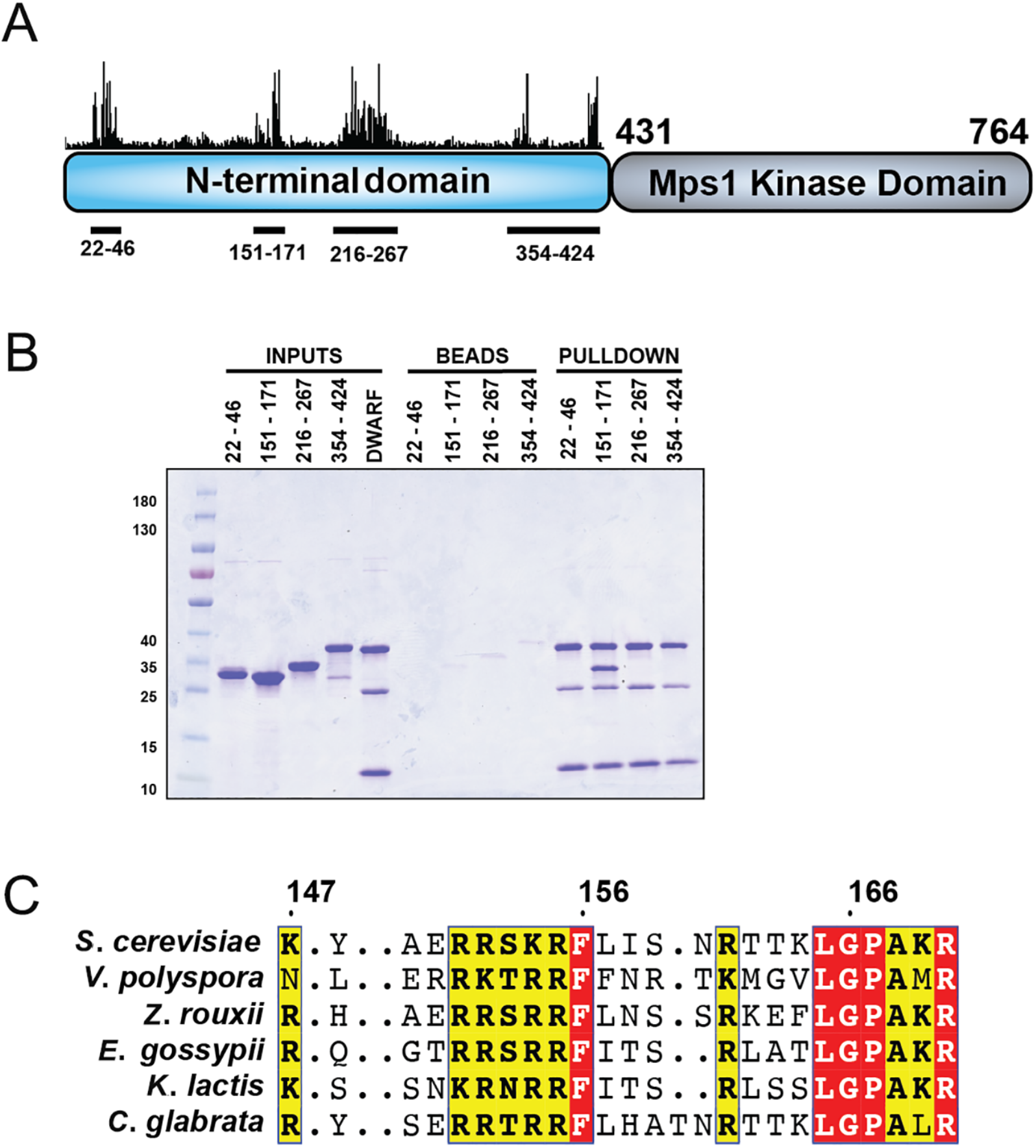
Binding of Mps1 N-terminal extension, residues 151–171, with Ndc80c^dwarf^. **(A)** Schematic representation of the domain structure of *S. cerevisiae* Mps1. The bar graph above the N-terminal domain represents conservation determined by iterative searches with Jackhmmer; these conserved stretches span the residues shown below the schematic. **(B)** Pulldown experiment testing whether each of the four conserved stretches in **(A)** bind Ndc80c^dwarf^. Ndc80c^dwarf^ was immobilized to saturation on Ni-NTA agarose and incubated with GST-fusions of each of the four conserved stretches. **(C)** Multiple sequence alignment showing conservation of residues 147-170 among point-centromere yeast.

### Binding of Mps1 and Mps1 mutant peptides to Ndc80c

We measured by fluorescence polarization the affinity of Mps1 for a peptide comprising residues 137-171. We labeled the peptide at its N-terminus with the Bodipy fluorophore and titrated Ndc80c^dwarf^ (Fig. 2A). We then used unlabeled peptide and peptide mutants in a competition format, at a fixed dwarf Ndc80c concentration (Figs. 2B,C). The mutants correspond to mutations characterized in one of the accompanying papers [40]. The Mps1 peptide displaced the fluorescent Mps1 peptide, at a concentration consistent with the affinity measured by direct titration. Mutating either the basic residues or the hydrophobic residues in the first part of the consensus motif or the conserved arginine at the end of the second part lowered affinity (raised the Ki) by roughly two logs; mutating both the final arginine and the hydrophobic residues eliminated any detectable competition altogether.

**Figure 2.**
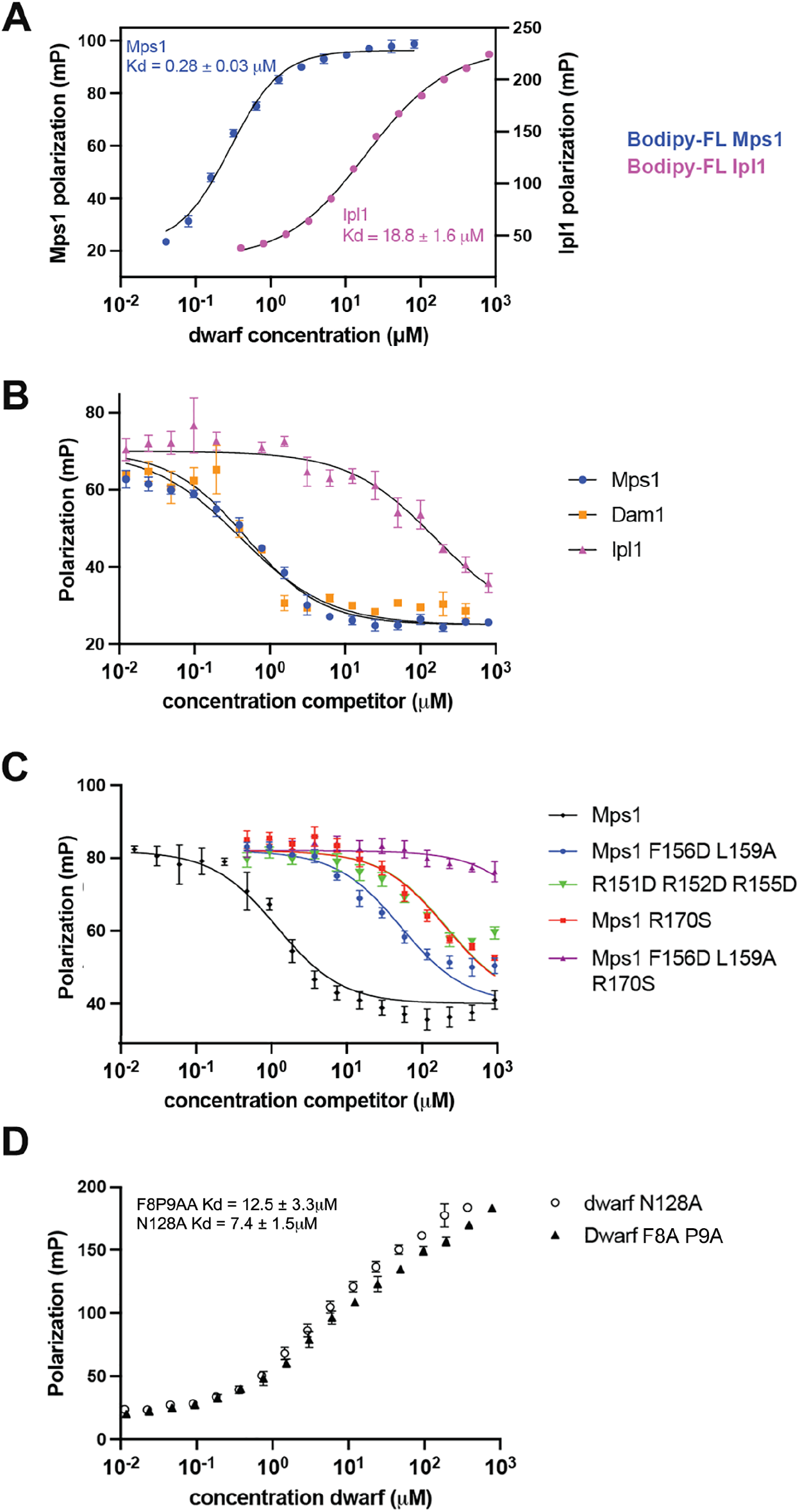
Fluorescence polarization measurements of peptides from Mps1 and Ipl1 binding with Ndc80c^dwarf^ and competition with each other and and with a Dam1 peptide. **(A)** Plot of fluorescence polarization from either 20 nM Bodipy-FL labelled Mps1(131-171) or 20 nM Bodipy-FL labeled Ipl1(26-59) as function of increasing Ndc80c^dwarf^ concentration. **(B)** Competition binding experiment in which decreasing fluorescence polarization shows competition with Bodipy-FL labelled Mps1c by unlabeled Mps1, Ipl1, and Dam1 competitor peptides for binding Ndc80c^dwarf^. **(C)** Competition binding experiment identical to that described in **(B)**, except the competitor peptides contained mutations expected to decrease affinity [48]. **(D)** Plot of fluorescence polarization for 20 nM Bodipy-FL labelled Mps1 in the presence of increasing concentrations of mutant Ndc80c^dwarf^ (see [40]).

We also determined the binding of the Mps1 peptide with two Ndc80c^dwarf^ constructs bearing mutations characterized in one of the accompanying papers -- N128A and F8A P9A [48]. In both cases, the Mps1 peptide affinity was about 2 logs weaker than for the wild-type protein (Fig. 2D).

### Structure of Ndc80c chimera with Mps1 binding motif

We validated conclusions from the competition binding experiments and AF2 predictions by determining x-ray crystal structures of Ndc80c^dwarf^ associated with the corresponding peptide segments from Mps1 and Ipl1. We applied the strategy we used to determine the Dam1-Ndc80c interaction [47], by generating chimeras with the C-terminus of the Mps1 or Ipl1 peptide joined to the N-terminus of Nuf2 as a continuous polypeptide chain. Because Ndc80c^dwarf^, designed originally to show the structure of the four-chain junction between the Ndc80:Nuf2 and Spc24:Spc25 heterodimeric components, crystallizes readily [51], it is a useful scaffold for constructing these chimeric proteins.

The Mps1: Ndc80c^dwarf^ chimera crystallized in space group C2221 (a = 72.5 Å, b = 168.5 Å, c = 229.6 Å). We recorded diffraction data to a minimum Bragg spacing of 2.9 Å and determined the structure as described in Methods (Fig. 3A; Table S1). Conservation of residues in the aligned Mps1 sequences for point-centromere yeasts indicates a bipartite motif (Fig. 1C): two conserved six-residue segments separated by 7 or 8 more variable residues. Contacts in the structure are consistent with these conservations and with the conservation of residues in Ndc80:Nuf2 with which the two six-residue segments interact. Fig.S1B shows a comparison of the AF2 prediction and the crystal structure.

**Figure 3.**
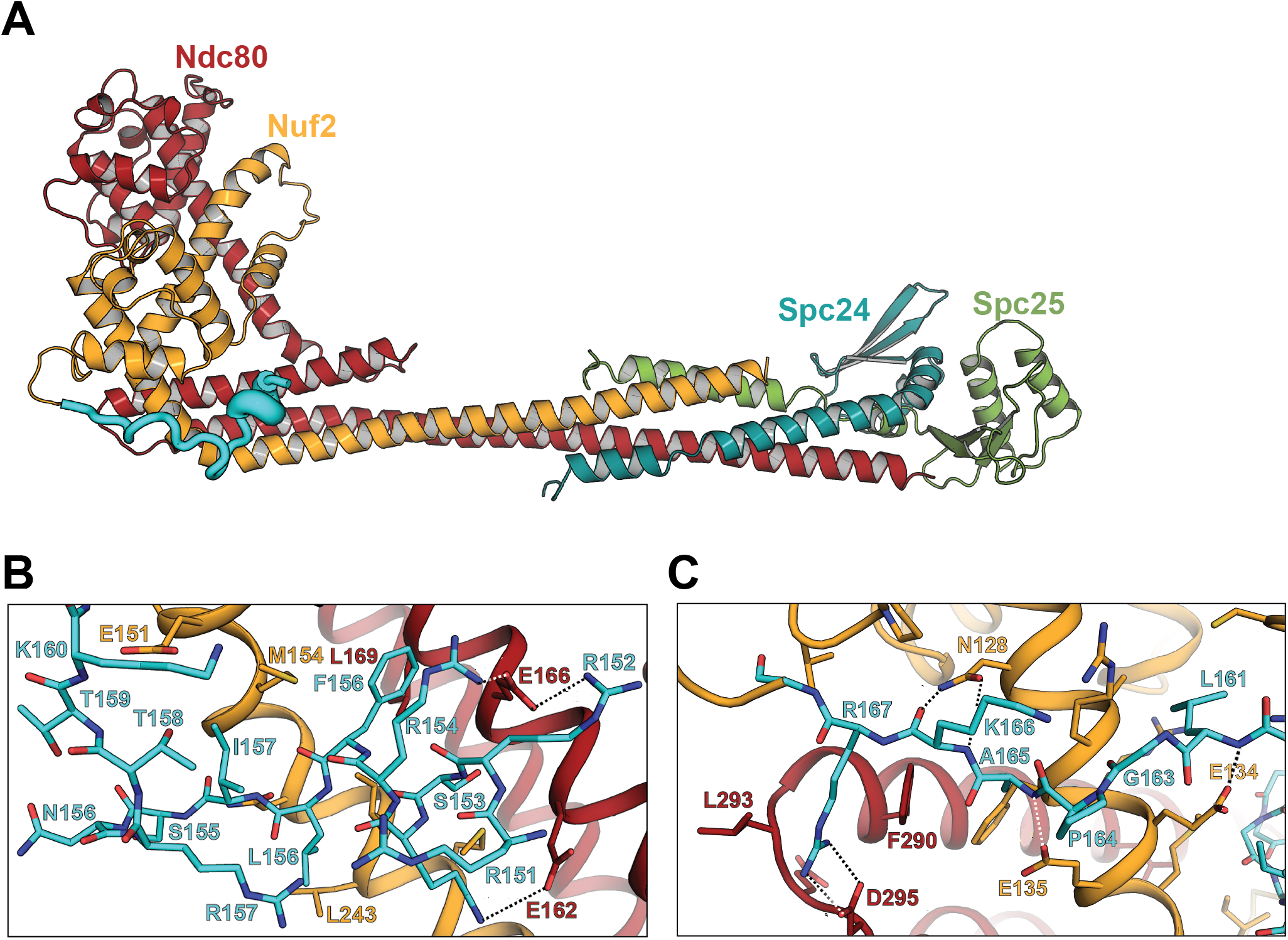
Crystal structure of Ndc80c^dwarf^ with Mps1 residues 124-176 were fused to the Nuf2 N-terminus. **(A)** Cartoon representation of the Ndc80c^dwarf^ heterotetramer with the Mps1-Nuf2 fusion. **(B)** Close-up view of the helical N-terminal portion of the Mps1 segment (rotated about 60° clockwise from the view in **A**). (**C)** Close-up view of the C-terminal portion of the Mps1 segment (oriented approximately as in **A**).

The N-terminal six residues of the motif (RRSKRF) form a short helix, which has a set of contacts with conserved glutamate residues in αH of Ndc80 and with conserved features of Nuf2 (Fig. 3B). In the crystal structure, the guanidinium groups of R152 and R154 both bridge to the carboxylate of Ndc80 E166, and the side-chain NH3+ of K153 projects to the carboxylate of Ndc80 E162. The conserved phenylalanine, Mps1 F156, inserts into a hydrophobic pocket lined by Ndc80 L169, L170 and F172, and Nuf2 M154, with a further contribution from Mps1 L157 (Fig. 3B).

The C-terminal six residues (LGPAKR) of the Mps1 motif extend across Nuf2 αG, with contacts from residues 128 and 130-132 (conserved in point-centromere yeast) (Fig. 3C). Asn 128, conserved not only in yeasts but also in metazoans, anchors the end of the motif with hydrogen bonds between its side-chain amide and the main-chain amide and carbonyl of K169 (the penultimate residue in the motif), thus mimicking the hydrogen bonds in an antiparallel β-sheet. This interaction positions the invariant arginine of the motif to salt bridge with Ndc80 Asp295, likewise conserved in both metazoans and yeast.

The structure rationalizes the phenotypes of mutants studied in the two accompanying papers, several of which we have incorporated into the binding measurements reported above. Our description of contacts with the N-terminal part of the bipartite motif can account for the effects of Mps1 mutations of basic and hydrophobic residues described in Pleuger et al [40] (designated there as br and hm mutants, respectively). They have also examined phosphomimetic and alanine mutations at S159, T162, and T163, all of which are at the Mps1:Nuf2 interface, in the loop between the two parts of the full motif. We expect that phosphorylation of T162 in particular, which faces a hydrophobic surface on Nuf2, at the junction between helices G and H, would displace that part of the bound peptide and presumably reduce affinity, but a displaced conformation would probably still allow the key residues in the binding motif to dock correctly.

Particular strong phenotypes among the Nuf2 mutations studies by Parnell et al [48] are at conserved residues F8P9 (mutated together), S124, and N128. We have described above the role of N128 in buttressing the bound peptide; it has the same key role in binding Dam1 [47]. Residues F8 and P9 together bury the hydrogen-bonding interactions of N128 with the main chain of the penultimate residue in the Mps1 and Dam1 binding motifs and probably contribute both to the strength of those interactions and to correct positioning of the invariant arginine to salt bridge with Ndc80 D295. Phosphorylation of Nuf2 Ser124 would displace F8 and generally reposition the Nuf2 N-terminal loop.

### Ipl1 interaction with Ndc80c

Ipl1 and Mps1 both phosphorylate residues in the N-terminal extension of Ndc80 [7, 8, 34, 53]. Documented Ipl1 kinetochore contacts are between its CPC partners, Sli15/INCENP and Bir1/survivin, and the inner-kinetochore COMA complex and the CBF3 component Ndc10, respectively [54–56]. In the CPC, a long, largely flexible segment (~600 residues) of Sli15/INCENP intervenes between the N-terminal region that interacts with Bir1 and Nbl1 and the segment that interacts with the Ipl1 kinase domain. There is also an approximately 100-residue, N-terminal extension of the Ipl1 kinase domain itself (Fig. 4A). We examined whether Ipl1 might attach directly to Ndc80:Nuf2 through contacts in this region by carrying out an AF2 prediction similar to the ones we had run for Mps1. Although the per-residue confidence level was much lower than for the Mps1 sequence (Fig. S1C), the interactions of the Ipl1 peptide were consistent from structure to structure among the 25 “top” predictions ouput by the program. Moreover, the key motif, SKIPSP(V/I)R, is present with slight variation, in Ipl1/AuroraB from other point-centromere yeast (Fig. 4B). The prediction suggested close overlap with the Mps1 interaction, with the final arginine of the motif conserved among point centromere yeast in both proteins. S. cerevisiae Ipl1 has two copies of the motif, while many other point-centromere yeasts have just one. The AF2 predictions split relatively evenly between the two alternatives in the S. cerevisiae sequence.

**Figure 4.**
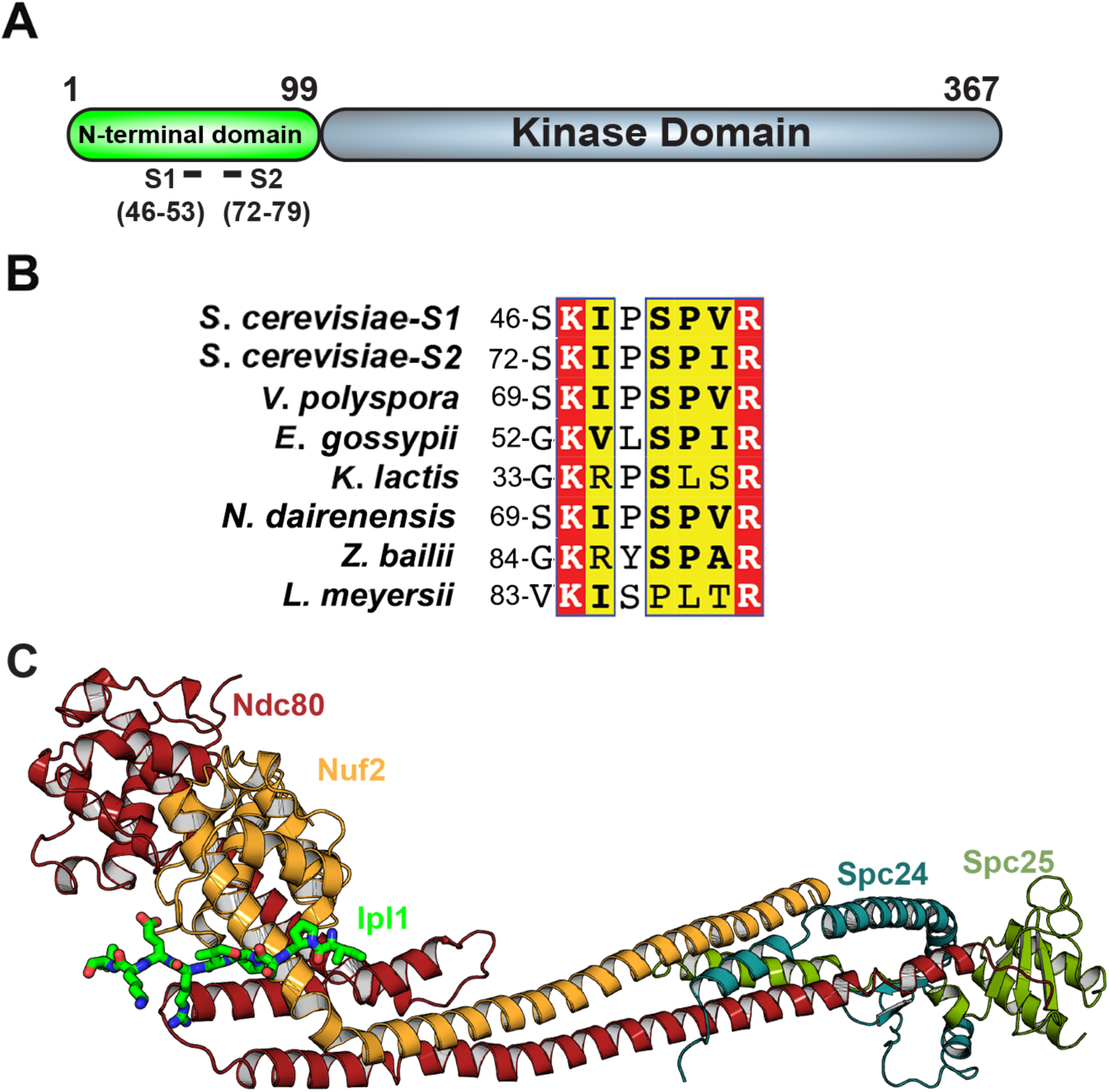
Crystal structure of a segment of the Ipl1 N-terminal domain that binds Ndc80c. **(A)** Schematic representation of the domain structure of Ipl1. “S1” and “S2” indicate each instance of a duplicated sequence that binds Ndc80c. **(B)** The sequences of S1 and S2 from **(A)** aligned with homologous segments of Ipl1 from other point-centromere yeast. **(C)** Cartoon representation of the crystal structure of Ndc80c^dwarf^ with Ipl1 S1 (residues 45-59) fused to the N-terminus of Nuf2.

We measured the Ipl1 peptide affinity, both directly and comparatively, with competition results that strongly supported site-overlap for all three (Fig. 2). The Ipl1 peptide bound more weakly than the other two, but avidity contributed by the two copies of the conserved motif would likely enhance binding of the intact protein.

### Structure of Ndc80c chimera with Ipl1 binding motif

We generated a chimeric construct that fused Ipl1 residues 45-59 to the N-terminus of Ndc80c^dwarf^ Nuf2. The chimera crystallized in space group P43212 (a=b=115Å,c=424Å), with two complexes in the asymmetric unit related by a non-crystallographic dyad. We recorded anisotropic diffraction data to a minimum Bragg spacing of 3.9Å, but intensities of reflections in the a* and b* directions fell below significance at Bragg spacings of ~6 Å. Despite the poorly ordered crystals, we determined phases by molecular replacement and calculated an electron density map at a nominal resolution of 6 Å (see Methods). When phased only on the molecular-replacement model, a difference map showed strong density for ordered parts of the Ipl1 segment and a structure overall that was consistent with the AF2 prediction between Ipl1 residues 48 and 56 (Fig. 4, Fig. S1D). The density for this segment was continuous with that for the Nuf2 subunit of the non-crystallographic-symmetry related complex (Fig. S2). Thus, the asymmetric unit is a “domain-swapped” dimer, and the interactions may have been essential for the crystals to form. Formation of the dimer by contacts in trans shows that binding of the Ipl1 segment in the location detected was not an artificial consequence of having fused it to the N-terminus of Nuf2.

### Comparison of Mps1, Ipl1 and Dam1 interactions with Ndc80c

Dam1 and Mps1 interact with the head of Ndc80c through segments in extended regions of their polypeptide chains. The competition experiment with the Dam1 peptide indicates that the Mps1 and Dam1 segments have very similar affinities for Ndc80c (Fig. 2B). The consensus sequence motif in both cases has two distinct parts, with a linker of variable length between them. The first part of the motif includes two or more basic residues followed by two hydrophobic residues; the second part, a tetrapeptide terminating in an invariant arginine. The contacts on Ndc80c are likewise conserved. The basic residues at the N-terminal end of the bipartite motif pair with conserved glutamic-acid residues on Ndc80 (Glu 269, 272, 276, *S. cerevisiae* numbering); the specific pairing may vary from ortholog to ortholog, as both arginine and glutamic-acid side chains can adopt multiple configurations. The pair of hydrophobic residues fits into a pocket between Nuf2 helices A and G. The invariant arginine at the end of the motif pairs though a salt bridge with Asp295 (*S. cerevisiae* numbering), likewise invariant (or in a few cases, glutamic acid) among point-centromere yeast. In so doing, it reaches across and caps the C-terminus of Ndc80 helix H. The interaction of Asn128 (*S. cerevisiae* numbering) with the main-chain hydrogen-bonding groups of the residues preceding the arginine is likewise universal, even in metazoans [38]. The conformations of the Mps1 and Dam1 peptides three residues to either side of the invariant arginine are the same, within the likely accuracy of the structure in that region and the presence of nearby crystal lattice contacts.

The Ipl1 consensus includes the essential features of the second part of the bipartite motif, but not those of the first. Moreover, the map, even though restricted to Bragg spacings greater than 6 Å, did not show clear density for polypeptide chain N-terminal to Ile 48 (or Ile 74, if fit with the other copy of the motif). The conformation of the peptide between residues 50 and 56 (or 76 and 82) appears to be very similar to that of the Mps1 motif, and hence also to the Dam1 peptide, even though the starting point for fitting the peptide was independent of the Mps1 structure.

The equilibrium dissociation constants of peptides for dwarf Ndc80c, as measured by direct titration with the relevant fluorescent peptide for Mps1 and Ipl1, are 280 nM and 18.8 μM, respectively (Fig. 2A). Their ratio is similar to the ratio (to wild-type) of the affinities of the Mps1 peptides in which residues in the first part of the bipartite motif have been mutated (Fig. 2C), consistent with the absence of the first part of the motif in Ipl1 and with the similarity of its motif with the second part of the bipartite Mps1 and Dam1 motifs.

## Discussion

The results presented above show that peptide segments in the flexible, N-terminal extensions of both Mps1 and Ipl1 bind a site on the head of Ndc80c that is essentially the same as the site bound by a segment in the C-terminal extension of Dam1. Ndc80c connects the centromere-proximal components of a kinetochore with the MTs of the mitotic spindle, and the DASH/Dam1c coordinates multiple Ndc80 complexes, thereby enabling end-on kinetochore-MT attachment. The Ndc80c head is thus a communication hub for ensuring a robust linkage between centromere and spindle and for integrating the SAC with tension sensing and error correction.

Mps1 is the sole co-purifying kinase on isolated, unattached kinetochores [34] [17]. We can infer that it is probably bound with Ndc80c through the contact described here, although there could be additional, so far undetected, interactions with segments elsewhere in the long N-terminal extension. Side-on attachment need not displace Mps1, and even end-on attachment, which requires the Dam1c/DASH, need not do so, as it includes contacts between Ndc80 and both Ask1 and Spc19/Spc34. Moreover, there are approximately 8 copies of Ndc80c in prometaphase/metaphase [32], and 16-17 copies of Dam1 in a single Dam1c/DASH ring [31]. Thus, even for the single kinetochore-microtubule attachment found in point-centromere yeast, mixed states are likely, in which Mps1 continues to be present (at decreasing levels) and hence continues to activate the SAC, until fully displaced by (dephosphorylated) Dam1. Autophosphorylation of Mps1 appears to result in its release (in vitro) from kinetochore association [39], but the relevant sites have not been determined. There are no serines or threonines in conserved positions within the bipartite motif, ruling out a direct regulation by phosphorylation of the Ndc80c interacting segment analyzed here.

In human Mps1, two distinct regions have been implicated in kinetochore interaction: a short N-terminal segment preceding a three-repeat TPR domain (features absent in yeast Mps1) and a conserved segment in the long “middle region” that links the TPR motifs with the kinase domain at the C-terminus [38]. Mutation of residues on the surface of the Ndc80 CH domain appear to affect binding of the former; mutation of N126 (corresponding to N128 in *S. cerevisiae*) to alanine interferes with binding of the latter. The sequence of the conserved, Mps1 middle-region segment includes a QSCPFGR motif, with invariant arginine in various vertebrate species; AF2 yields very high confidence binding of that segment (and several residues C-terminal to it) at the site in the human Ndc80c head corresponding precisely to the communication hub site we have characterized in yeast (Fig. S4A). The Nuf2 interaction thus appears to be present in metazoans as well as in yeast.

Several lines of evidence suggest that Mps1 may under some circumstances recruit DASH/Dam1c to kinetochores, by modification of residues on Dam1 or on another component of the heterodecameric complex [48, 57]. Principal candidates are Dam1 residues Ser218 and Ser 221, which are part of a conserved motif in point-centromere yeast Dam1 (218-S/TxASFVxNP-226, *S. cerevisiae* numbering, preceded by several negatively charged residues)[58]. Available structural information does not provide hypotheses about possible binding partners or about autoinhibition mechanisms for ring formation that might involve this short, conserved motif, although conservation suggests a defined function. The other reported Mps1 site in Dam1, Thr15, is part of the N-terminal “staple” that connects one DASH protomer with another; phosphorylation of Thr15 by Mps1, like that of Ser20 by Ipl1, might therefore disfavor ring formation, rather than stabilize it [59].

Ipl1/Aurora B has at least two distinct roles in regulating kinetochore assembly and attachment. Phosphorylation of Dsn1 activates the MIND/Mis12 complex by relieving autoinhibition and allowing binding of Mif2/CENP-C and Ame1 to sites on the Mtw1-Nnf1 globular “head [60, 61] [62, 63]. Subsequently, the kinase mediates error correction and reverses end-on attachment in the absence of tension by phosphorylating both the Ndc80 N-terminal extension and at least three components of DASH/Dam1c -- Dam1 itself, Ask1, and Spc34 [33] -- or the Ska complex in metazoans [64]. In yeast, binding of Bir1 with Ndc10 can recruit Ipl1 early in kinetochore assembly, which begins immediately after replication of yeast centromeres in early S phase, and the Ndc10-Bir1 interaction might facilitate access to Dsn1. Binding of Sli15 with COMA appears to provide a further anchor point, once the Ctf19 complex (Ctf19c) has assembled on the Cse4 nucleosome [54, 55]. The interaction with the Ndc80c head could in principle then localize the kinase domain close to its targets. About 600 Si15 residues intervene between its attachments to Bir1 and Ipl1; these could readily stretch between CDEIII, to which Ndc10 binds, and the head of Ndc80, or to Dam1 after end-on attachment, but whether Ndc80c head binding by the N-terminal extension of Ipl1 can occur concomitantly with the Sli15-COMA interaction will depend on the position, not yet determed, of the relevant residues on Sli15. The proximity of the Ipl1 kinase domain afforded by interaction between the Ndc80c head and the N-terminal extension might be important for capturing target sites on the Ndc80 N-terminal extension when some or all of those residues dissociate, in the absence of tension, from binding with a microtubule. Likewise, when tension is absent after end-on attachment, the same tethering might enhance phosphorylation of the motif close to the C-terminus of Dam1.

The Ipl1 motif that binds Ndc80c is also the target of phosphorylation by Cdk1, which preferentially modifies serine or threonine at sites with the sequence (S/T)Px(R/K) [65]. The current models (crystallographic or AF2 predicted) do not clearly suggest whether phosphorylation of the targeted serines (S50 and S76) would strengthen or weaken Ipl1 binding, as the serines lie between conserved arginine and glutamic acid residues on Nuf2. A Bim1 binding motif SxIP [66] immediately precedes the SP(V/I)R Ndc80c-binding sequences in *S. cerevisiae* Ipl1 (Fig. 4B), and phosphorylation of S50 and S76 appears to suppress Ipl1 association with Bim1 until activity of the Cdk1 kinase Cdc28 declines in anaphase [65]. Although the Bim1-binding consensus is absent in many other point-centromere yeast, the Cdk1 consensus is almost always present.

Human Aurora B also has a candidate Ndc80c binding motif (TLPQR) in the 70-residue N-terminal extension that precedes the kinase domain in the polypeptide chain. AF2 predicts with high confidence an association very similar to the predicted Mps1 association, including interaction of several residues C-terminal to the key arginine (Fig. S4B). Thus, as with Mps1, the Ndc80c head is probably also an important communication hub in metazoans.

Detection of a segment in the Ipl1 N-terminal extension that interacts with the same site on the Ndc80c head that binds Dam1 and Mps1 and observations that Ipl1 and Mps1 have partly overlapping roles in error correction raise the question of when, in the interval from S-phase to late prometaphase, does each of these kinases have its most important function and what are its critical targets. Ipl1 appears to be required for error correction throughout prometaphase, even after formation of two distinct kinetochore clusters, which can still contain syntelic attachments [21]. Mps1 must likewise be present, at least on any pair not under tension, to keep the SAC engaged. Future experiments will need to determine the trade-off among Mps1, Ipl1, and Dam1 at the Ndc80c communication hub, the respective phosphorylation targets when Mps1 or Ipl1 binds there, and the importance of mixed states, with some of the Ndc80 complexes on any kinetochore bearing one of the kinases, some bearing the other, and even some bound with Dam1.

## Methods

### Protein purification

The constructs used to express Ndc80c^dwarf^ and its variants are identical to those used previously [51], except for the exclusion of a non-native sequence N-terminal to Nuf2 that promoted crystallization in the earlier study and the addition of Mps1 and Ipl1 N-terminal extensions as described below. The purification scheme was identical for all variants, except for the final gel-filtration step, for which the buffer composition varied depending on the subsequent application. Briefly, shortened versions of Ndc80/Nuf2 and Spc24/Spc25 were cloned into the orthogonal expression vectors pETduet1 and pRSFduet, respectively. Proteins were co-expressed in Bl21 (DE3) *E. coli* (New England Biolabs). Cells were grown at 37°C in 2XYT media to an OD600 of 0.8, induced with 500 µM IPTG, and incubated overnight with shaking at 225 rpm at 18°C. Cell pellets from 6 L of culture were resuspended in 100 mL of buffer containing 100 mM Tris pH 8.0, 250 mM NaCl, 10 mM imidazole pH 8.0, 2 mM tris(2-carboxyethyl)phosphine (TCEP), 1 mM PMSF, 2 “Complete” protease inhibitor tablets (Roche), 25 μg/mL lysozyme (Gold Biochem) and 5 μg/mL DnaseI (Gold Biochem). Cells were lysed by sonication and the lysate clarified by centrifugation and applied to Ni-NTA agarose equilibrated with 20 mM Tris pH 8.0, 100 mM NaCl, 10 mM imidazole pH 8.0, 2 mM TCEP (equilibration buffer). Bound material was washed with equilibration buffer containing 20 mM imidazole and 500 mM NaCl and then with equilibration buffer. Proteins were eluted with equilibration buffer containing 400 mM imidazole pH 8.0 and 100 mM NaCl. Eluates were treated overnight with TEV protease to remove the 6-His tag from the N-terminus of Ndc80 (except for protein to be immobilized for pulldown experiments) and subjected to anion exchange chromatography using a Hi-trap Q HP (Cytiva), followed by size-exclusion chromatography using a Hi-load Superdex 200 16/60 column (Cytiva). For protein crystallization the column was equilibrated with 10 mM HEPES pH 7.2, 100 mM NaCl, and 2 mM TCEP. For fluorescence polarization binding and pulldown experiments, the buffer was 10 mM HEPES pH 7.2, 150 mM NaCl, 2 mM TCEP.

The Mps1 fragments used for pulldown experiments were ordered from Genscript and cloned into pGEX6P1 (Novagen) to express them as GST fusions. Cells were grown at 37°C in 2XYT media to an OD600 of 0.8, induced with 500 µM IPTG, and incubated overnight with shaking at 225 rpm at 18°C. Cell pellets from 2 L of culture were resuspended in 30 mL of buffer containing 100 mM Tris pH 8.0, 250 mM NaCl, 2 mM TCEP, 1 mM PMSF, 25 mg/mL lysozyme (Gold Biochem), 5 mg/mL DNaseI (Gold Biochem) and a single “Complete” protease inhibitor tablet (Roche). Cells were lysed by sonication and the lysate clarified by centrifugation and applied to glutathione agarose (1 mL bed volume) (Pierce). The resin was washed with 20 mM Hepes pH 7.2, 250 mM NaCl, and 2 mM TCEP (wash buffer). Protein was eluted from the resin by applying 6 mL of wash buffer supplemented with 20 mM reduced glutathione (Sigma).

### Pulldown Experiments

For the pulldown experiments in Figure 1, Ndc80c^dwarf^ with a 6-His affinity tag on the N-terminus of Ndc80 was immobilized to saturation on Ni-NTA agarose by incubation with gentle agitation at 4°C. Beads were pelleted by centrifugation, washed three times in 20 mM HEPES pH 7.2, 150 mM NaCl, 2 mM imidazole pH 7.2, 2 mM TCEP. Next, the beads were incubated with the Mps1-GST fusion eluted from the glutathione agarose for 30 min. Beads were again pelleted, washed three times with wash buffer, pelleted a final time, and the wash buffer aspirated from the tube. To elute bound proteins, 50 µL of wash buffer supplemented with 500 mM imidazole pH 7.2 was added to ~25 µL of beads. After an additional spin, the eluted proteins were visualized by SDS-PAGE.

### Peptide binding experiments

Mps1 (137-171), Ipl1 (26-59), and Dam1 (254-305) peptides with C-terminal cysteines, were synthesized by the Tufts University Core Facility. The Mps1 peptide was labelled with Bodipy FL maleimide (Thermo Fisher) according to the manufacturer’s instructions. The labeled peptide was separated from unreacted dye by cation exchange chromatography with Source 15S resin (Cytiva). Fluorescence polarization was measured at 25°C with an Envision plate reader (Perkin Elmer) equipped with a filter set optimized for fluorescence polarization of FITC, which has the same absorption and emission spectra as those of Bodipy FL. For direct binding experiments, an 860 μM solution of Ndc80c^dwarf^ in a buffer containing 20 mM Hepes pH 7.2, 150 mM NaCl, 2 mM TCEP was subject to 2-fold serial dilutions and subsequently supplemented with an equal volume of a buffer-matched solution of 40 nM Bodipy-FL labelled Mps1 peptide. Fluorescence polarization was measured in triplicate. For competition binding experiments, 2 mM solutions of each peptide, dissolved in 20 mM tris pH 7.2, 100 mM NaCl, 2 mM TCEP, was subject to serial 2-fold dilutions. An equal volume of a buffer-matched solution containing 40 nM Bodipy-FL and 1 μM Ndc80c^dwarf^ was added to each of the dilutions, and fluorescence polarization was measured in triplicate.

### Ndc80c^dwarf^-Mps1 fusion crystallization and structure determination

Ndc80c^dwarf^, with residues 124-176 of *S.cerevisiae* Mps1 appended to the N-terminus of Nuf2, was concentrated to 25 mg/mL in 10 mM Hepes pH 7.2, 100 mM NaCl, 2 mM TCEP. Crystals were grown by vapor diffusion in hanging drop format in 24-well plates (Hampton Research). Crystals used for diffraction data collection grew from drops in which 1 μL of protein solution was mixed with 1 μL of a well solution containing 13% polyethylene glycol 8000, 1M sodium chloride, 100 mM PIPES pH 6.1. Crystals were harvested, soaked for 1-2 minutes in a well solution supplemented with 25% glycerol, and immersed immediately in liquid nitrogen. The complex crystallized in space group C222_1_ (a = 72.52 Å, b = 168.45 Å, c = 229.57 Å). Data to a minimum Bragg spacing of 3.29 Å were recorded on beamline 201 at the Advanced Light Source, indexed, integrated, using DIALS, as implemented in xia2 [67]. The data were scaled and merged using the STARANISO server [68]. The structure was determined by molecular replacement in Phenix [69], using three non-overlapping segments of Ndc80c^dwarf^ (PDB 5TCS) as search models. The molecular-replacement maps showed clear density for the Mps1-derived peptide. Model building was carried out in Coot [70] and refinement, in Phenix [69]. Data statistics are in Table S1.

### Dwarf Ipl1 fusion crystallization and structure determination

Ndc80c^dwarf^, with residues 45-59 of *S.cerevisiae* Ipl1 appended to the N-terminus of Nuf2 was concentrated to 40 mg/mL in 10 mM Hepes pH 7.2, 100 mM sodium chloride, 2 mM TCEP. Crystallization screens were set up in sitting drop format using a Mosquito liquid handling robot. The 400 nL drops contained equal volumes of protein and well solution. Crystals grew from drops equilibrated with 15% polyethylene glycol 2000 monomethyl ether, 500 mM sodium chloride, 50 mM Tris pH 7.5. Crystals were harvested directly from the screening drop, soaked for 1-2 minutes in a well solution supplemented with 25% glycerol, and immersed immediately in liquid nitrogen. The complex crystallized in space group P4_3_2_1_2 (a = 114.81 Å, b = 114.81 Å, c = 424.17 Å). Data were recorded on beamline 201 at the Advanced Light Source and integrated with XDS [71]. Because of substantial anisotropy in the diffraction, data were scaled and merged using the STARANISO server [68]. The minimum Bragg spacing following anisotropic scaling was 3.93 Å in the “best” direction and 6.76 Å in the “worst”. We determined the structure by molecular replacement in Phenix [69], using three search models derived from Ndc80c^dwarf^ (PDB 5TCS). There were two complexes in the asymmetric unit. The 2Fo-Fc and Fo-Fc molecular replacement maps had clear density for the appended Ipl1 peptide in both non-crystallographic symmetry (ncs) related copies. The density also showed that the peptide appended to one ncs copy of Ndc80c^dwarf^ was bound to the ncs-related copy -- i.e., that the two ncs-related complexes are a “domain-swapped” pair (Fig. S2). We used O [72] to fit a peptide comprising residues 48-56 of Ipl1 into the Fo-Fc density, truncated to 6 Å in all directions, using the AF2 model as a starting point. Refinement was carried out in Phenix [69]. Statistics are in Table S1.

### Data availability

Coordinates and structure factors for the two structures reported here have been deposited in the Protein Databank with accession numbers 8V10 and 8V11.

## Acknowledgments

We are grateful to Matthew Miller and Emily Parnell for careful reading and commenting on the manuscript, and to Stefan Westermann, Richard Pleuger, Matthew Miller, Emily Parnell and Erin Jensen for discussions and for communicating results before submission of their papers. We thank the staff at Advanced Photon Source Beamline 201 for assistance with x-ray data collection. The research was supported by the Howard Hughes Medical Institute (SCH Investigator’s budget) and by Postdoctoral Fellowship PF-21-188-01-CCB to JAZ from the American Cancer Society.

## Figure Legends

**Supplemental Figure S1.** Comparison of the Mps1 and Ipl1 x-ray structures with their respective AF2 predictions. **(A)** AF2 multimer prediction of the Ndc80/Nuf2 CH domains and Mps1 residues 131-180. The pLLDT score is indicated by both color and diameter of the “cartoon putty”. **(B)** Superposition of the AF2 prediction of Mps1-bound Ndc80c^dwarf^ and the corresponding x-ray structure. **(C)** AF2 multimer prediction of the Ndc80/Nuf2 CH domains and Ipl1 residues 26-59. **(D)** Superposition of the AF2 prediction of Ipl1-bound Ndc80c^dwarf^ and the corresponding x-ray structure.

**Supplemental Figure S2.** Domain swap in Ipl1 chimera with Ndc80c^dwarf^. Density at the interface between two non-crystallographic symmetry-related complexes shows that the Ipl1 peptide added to the N-terminus of Nuf2 in one complex associates with the Ndc80:Nuf2 head in the other. The Ipl1-Nuf2 in the right-hand complex is in light orange; the Ipl1-Nuf2 in the left-hand complex is in magenta; the magenta Ipl1-peptide associates with the orange Nuf2, and the orange Ipl1 peptide associates with the magenta Nuf2, with continuous density connecting across the (approximate) non-crystallographic twofold. The Ndc80 chains are in red and gray.

**Supplemental Figure S3.** AF2 predictions for human Mps1 and Ipl1 association with the head of human Ndc80c. **(A)** AF2 multimer prediction for the Ndc80/Nuf2 CH domains and segment from human Mps1 “middle region” (265-TKQSC**PFGR**VPVNLLNSPDCD-285; tetrapeptide terminating in invariant arginine in boldface). **(B)** AF2 multimer prediction for the Ndc80/Nuf2 CH domains and segment from the N-terminal extension of human Aurora B (20-GLST**LPQR**VLRKEPV-34: tetrapeptide terminating in invariant arginine in boldface).

