## Supplemental figures and tables for "A communication hub for phosphoregulation of kinetochore-microtubule attachment"

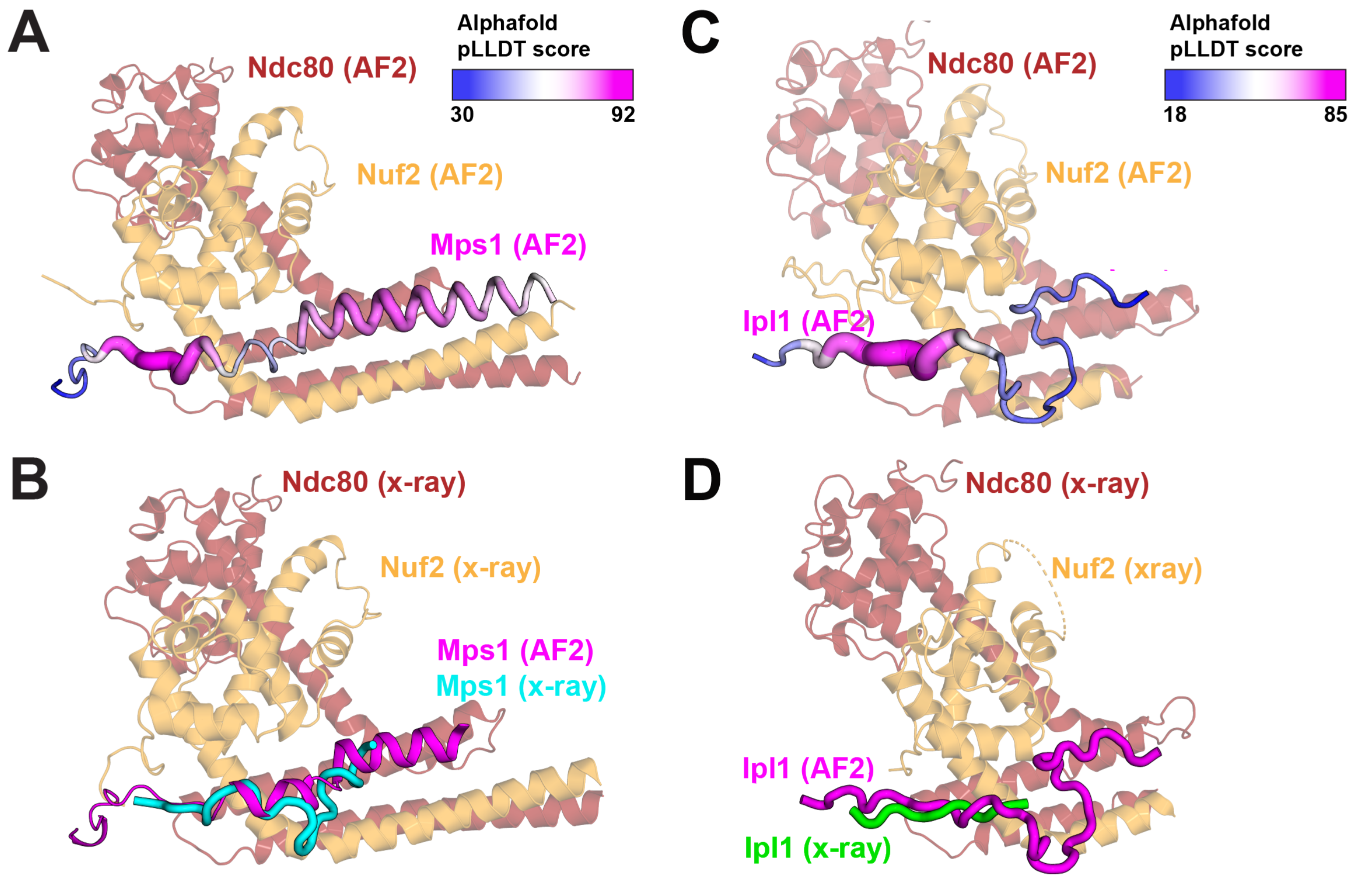


**Figure S1.** Comparison of the Mps1 and Ipl1 x-ray structures with their respective AF2 predictions. **(A)** AF2 multimer prediction of the Ndc80/Nuf2 CH domains and Mps1 residues 131-180. The pLLDT score is indicated by both color and diameter of the “cartoon putty”. **(B)** Superposition of the AF2 prediction of Mps1-bound Ndc80c$^{dwarf}$ and the corresponding x-ray structure. **(C)** AF2 multimer prediction of the Ndc80/Nuf2 CH domains and Ipl1 residues 26-59. **(D)** Superposition of the AF2 prediction of Ipl1-bound Ndc80c$^{dwarf}$ and the corresponding x-ray structure.

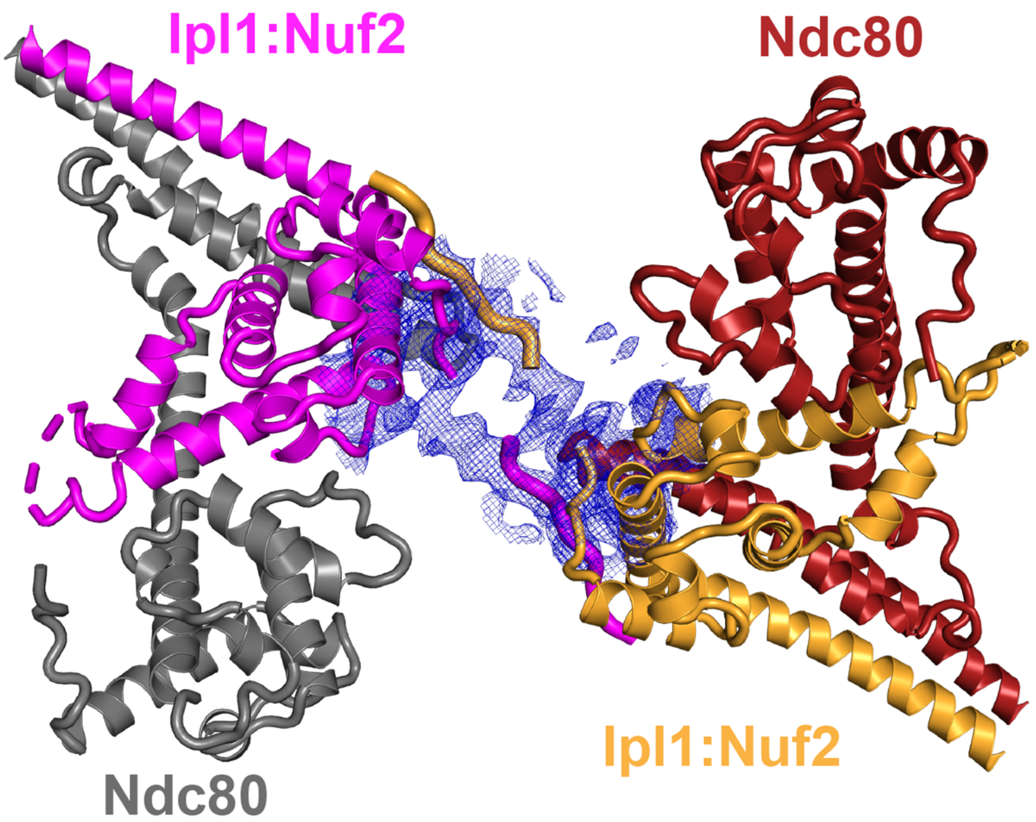


**Figure S2**. Domain swap in Ipl1 chimera with Ndc80c[dwarf]. Density at the interface between two non-crystallographic symmetry-related complexes shows that the Ipl1 peptide added to the N-terminus of Nuf2 in one complex associates with the Ndc80:Nuf2 head in the other. The Ipl1-Nuf2 in the right-hand complex is in light orange; the Ipl1-Nuf2 in the left-hand complex is in magenta; the magenta Ipl1-peptide associates with the orange Nuf2, and the orange Ipl1 peptide associates with the magenta Nuf2, with continuous density connecting across the (approximate) non-crystallographic twofold. The Ndc80 chains are in red and gray.

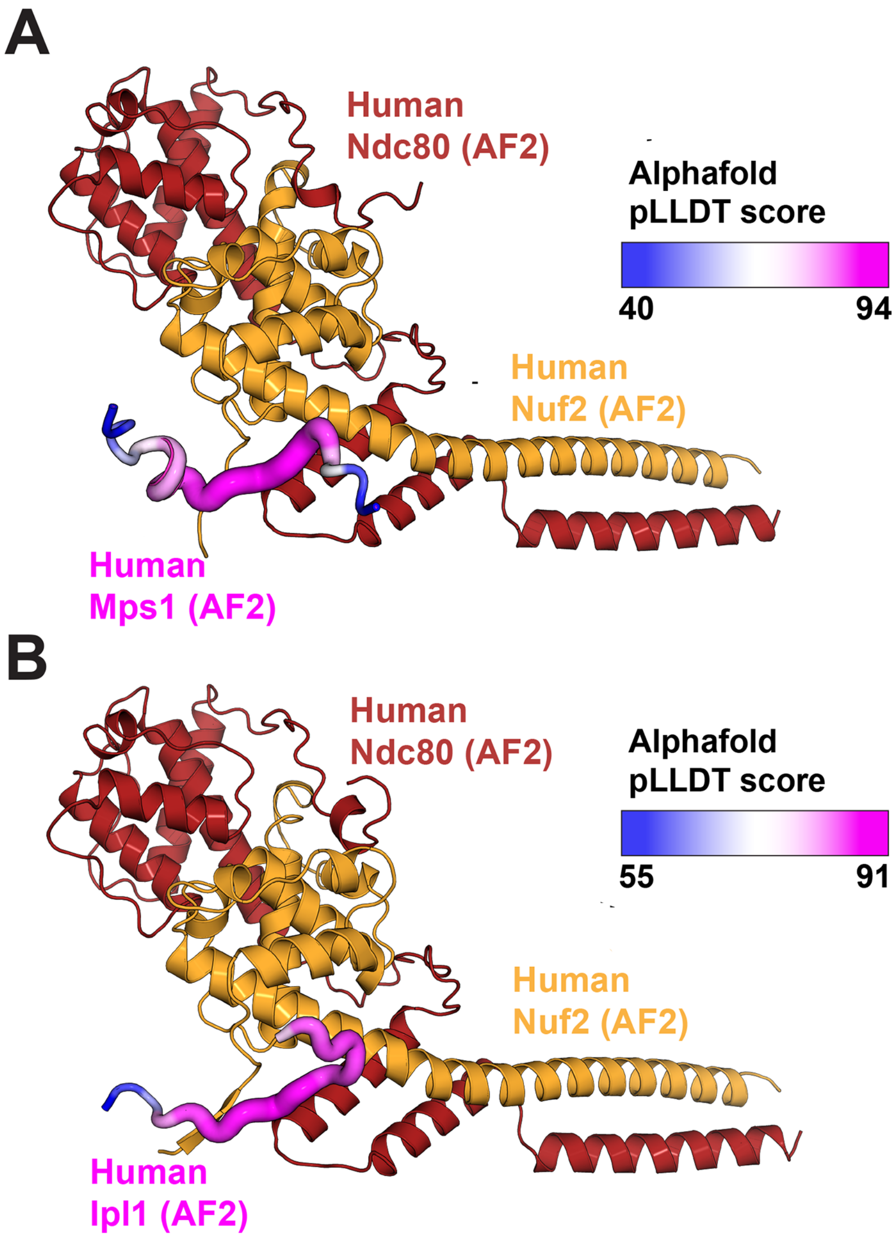


**Figure S3**. AF2 predictions for human Mps1 and Ipl1 association with the head of human Ndc80c. **(A)** AF2 multimer prediction for the Ndc80/Nuf2 CH domains and segment from human Mps1 "middle region" (265-TKQSC**PFGR**VPVNLLNSPDCD-285; tetrapeptide terminating in invariant arginine in boldface). **(B)** AF2 multimer prediction for the Ndc80/Nuf2 CH domains and segment from the N-terminal extension of human Aurora B (20-GLST**LPQR**VLRKEPV-34: tetrapeptide terminating in invariant arginine in boldface).

### Table S1. X-ray data collection and structure refinement statistics

| | **Ndc80:Mps1-Nuf2** | **Ndc80:Ipl1-Nuf2** |
|---|---|---|
| **Data collection** | | |
| Space group | $C222_1$ | $P4_32_12$ |
| Cell dimensions | | |
| a, b, c (Å) | 72.65 168.30 230.21 | 114.76, 114.76, 423.84 |
| α, β, γ (º) | 90.0 90.0 90.0 | 90.0, 90.0, 90 |
| STARANISO analysis | Diff. limit #1: 3.728, (1.0, 0.0, 0.0), **a*** | Diff. limit #1: 6.237, (1.0, 0.0, 0.0), **a*** |
| Diffraction limits (Å) [a] | Diff limit #2: 3.542, (0.0, 1.0, 0.0), **b*** | Diff. limit #2: 6.237, (0.0, 1.0, 0.0), **b*** |
| | Diff limit #3: 2.912, (0.0, 0.0, 1.0), **c*** | Diff. limit #3: 3.361, (0.0, 0.0, 1.0), **c*** |
| | Eigenvalue #1: 141.07, (1.0, 0.0, 0.0), **a*** | Eigenvalue #1: 107.1, (1.0, 0.0, 0.0), **a*** |
| Anisotropic tensor ($Å^2$) [b] | Eigenvalue#2: 141.37, (0.0, 1.0, 0.0), **b*** | Eigenvalue #2: 107.1, (0.0, 1.0, 0.0), **b*** |
| | Eigenvalue#3: 65.62, (0.0, 0.0, 1.0), **c*** | Eigenvalue #3: 48.19, (0.0, 0.0, 1.0), **c*** |
| Wavelength (Å) | 1.03094 | 1.0309 |
| Resolution (Å) [c] | 115.11–3.03 (3.4-3.03) | 48.53–3.95 (4.48–3.95) |
| $R_{merge}$ (%) | 15.5 (85.3) | 55.6 (390.0) |
| $R_{meas}$ (%) | 18.1 (103.9) | 57.3 (354.6) |
| $R_{pim}$ (%) | 0.68 (6.37) | 13.5 (123) |
| I/σI | 5.3 (1.6) | 6.7 (1.6) |
| $CC_{1/2}$ | 99.4 (68.3) | 99.3 (60.9) |
| Completeness spherical (%) | 64.9 (11.2) | 42.6 (10.5) |
| Completeness ellipsoidal(%) | 88.9 (48.1) | 89.0 (76.6) |
| Total number of observations | 111103 (3752) | 191124 (14902) |
| Unique reflections | 18,187 (909) | 10,981 (785) |
| Redundancy | 6.1 (4.1) | 17.4 (19) |
| **Refinement** | | |
| Resolution (Å) | 115.107–3.21 (3.41–3.21) | 48.53–3.95 (4.35–3.95) |
| No. of reflections used | 16929 (735) | 104112 (522) |
| Reflections used for R-free | 890 (39) | 559 (32) |
| $R_{work}$ / $R_{free}$ (%) | 24.19 (27.32) / 35.19 (38.64) | 30.19 / 34.83 (35.4 / 43.2) |
| $CC_{work}$ / $CC_{free}$ | 95.1 (0.5) / 0.911 (0.888) | 0.92 / 0.89 (0.30 / -0.30) |
| **Model statistics** | | |
| No. of non-hydrogen atoms | 5725 | 11,084 |
| B factors | | |
| Average <B> ($Å^2$) | 87.7 | 147.6 |
| $\langle B_i - B_j \rangle$ ($Å^2$) | 9.44 | 12 |
| R.m.s deviations | | |
| Bond lengths (Å) | 0.001 | 0.004 |
| Bond angles (º) | 0.35 | 0.65 |
| Rotamer outliers (%) | 0.15 | 0.1 |
| Ramachandran angles | | |
| Favored (%) | 96.36 | 96.21 |
| Outliers (%) | 0 | 0.68 |
| MolProbity clash score | 4.63 | 13.27 |
| PDB-ID | 8V10 | 8V11 |

[a] Diffraction limits (Å) and corresponding principal axes of the ellipsoid fitted to the diffraction cut-off surface as direction cosines in the orthogonal basis (standard PDB convention), and in terms of reciprocal unit-cell vectors.
[b] Eigenvalues of overall anisotropy tensor on |F|s ($Å^2$) and corresponding eigenvectors of the overall anisotropy tensor as direction cosines in the orthogonal basis (standard PDB convention), and in terms of reciprocal unit-cell vectors.
[c] Highest resolution shell is shown in parenthesis.
[d] After anisotropy correction and B-sharpening.
N.A.: not applicable.
